# Pseudo Dollo models for the evolution of binary characters along a tree

**DOI:** 10.1101/207571

**Authors:** Remco R. Bouckaert, Martine Robbeets

**Affiliations:** Centre of Computational Evolution, University of Auckland,, Auckland, New Zealand; Max Planck Institute for the Science of Human History, Jena, Germany

## Abstract

The stochastic Dollo model is a model for capturing evolution of features, for example cognate data in language evolution. However, it is rather sensitive to borrowing events, coding errors, semantic shift and other anomalies, so other models, in particular the covarion model, tends to have a better fit to the data. Here, we introduce the pseudo Dollo model, a model of character evolution along a tree that can be formulated as a three-state continuous time Markov chain (CTMC) model. The initial state represent absence of a feature, then a birth event allows the feature to be present. A death event can follow so that the feature becomes absent again. However, no new birth events are allowed after a death event has taken place.

We examine the model in a fully Bayesian setting, and demonstrate it can have a better fit than some of the popular alternative models on some real world datasets. Some variations on the pseudo Dollo model are introduced as well, including the multi-state pseudo Dollo model and pseudo Dollo covarion model.

The model is implemented in open source software Babel, a package to BEAST [2] licensed under LGPL. A user friendly way to set up an analysis is available through BEAUti, the graphical user interface of BEAST.

## 1 Background

The stochastic Dollo model [15] is a model of character evolution along a tree that is based on the Dollo principle [5]: a feature that is lost won’t be regained. Its corollary implies that a feature only evolves into presence once, but can be lost several times. This is a natural explanation of language evolution of cognates. After all, the definition of a cognate is a word form with a commonancestry. When there are few borrowing events in the cognate data, nor much coding errors, the stochastic Dollo model fits the data very well. However, when there are borrowing events across distant languages in the language tree, or there are coding errors, the stochastic Dollo model tends to have difficulty fitting the data, and two-state CTMC or covarion models [18] tend to fit better. Another situation where the stochastic Dollo model has problems is when a feature is modeled along a gene tree that suffers from incomplete lineage sorting and the feature of interest is the result of another cognate history than the one in the analysis. In this case, the feature may be Dollo like on the gene tree (that is, the tree representing the history of one of the meaning classes), but not necessarily on the species tree (that is, the history of languages). Semantic shift is another source of trouble for the stochastic Dollo model, where the word form remains present in a language, but its meaning changes between a number of related definitions.

The covarion model [18] in particular tends to show an unreasonable good fit in these cases. For example, when the data is mostly Dollo like, but also has a good number of borrowing events, like is the case for Indo-European [3, 11] and Austronesian [12] cognate data. It is hard to get an intuition why the covarion model works so well. Here, we introduce an alternative model to the stochastic Dollo model that fits well, and is intuitive like the stochastic Dollo model. As a bonus, a tree likelihood using the pseudo Dollo model can be calculated more efficiently than the Dollo model - the BEAST implementation is about twice as fast. We also introduce some variations on the pseudo Dollo model, and show it performs well on cognate data for Transeurasian languages.

## 2 Methods

First, we describe the binary CTMC, covarion and stochastic Dollo models, which are popular models for cognate evolution [3, 11, 12].

### 2.1 Two state CTMC model

A popular method for cognate evolution is the two state continuous time Markov chain (CTMC) model (see [17] for an introduction to CTMC models),which simply recognises two states: present and absent. The rate matrix is

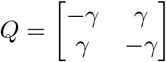

with non-negative frequencies *f*_0_, *f*_1_ such that *f*_0_ + *f*_1_ = 1. To guarantee identifiability, we set *γ* = 1, so there is just one degree of freedom left from the frequencies, which makes it the model with the fewest number of parameters among popular cognate evolution models. The model allows frequent switches between features being present and absent, unlike the stochastic Dollo model, so it is a lot less sensitive to borrowing and coding errors. When different sites evolve at different rates, the binary CTMC model can benefit from gamma rate heterogeneity across sites [19].

### 2.2 Covarion model

The covarion model [18] is a model that assumes a character can evolve in two different modes: a slow and a fast one. When in the fast mode, the character evolves quickly between absent and present states and vice versa, but when in the slow mode, it can stay in the same state for long periods of time. So, there are four states: fast absent, fast present, slow absent and slow present. When dealing with cognate data, absence of a cognate is encoded as both fast and slow absent, and presence of a cognate asboth fast and slow present, so it uses ambiguity codes. One way to encode the covarion model is through a Q matrix

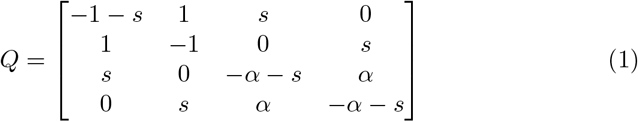

where *s* is the switch rate and *α* the mutation rate for the slow state. The fast mutation rate is fixed at 1 to ensure identifiability of the other parameters. Note this encoding is slightly non-standard, but tends to perform well on cognate data. Frequencies are parameterised through visible frequencies *f*_0_, *f*_1_ and hidden frequencies *h*_*fast*_, *h*_*slow*_, constrained such that *f*_0_ + *f*_1_ = 1 and *h_fast_* + *h_slow_* = 1 and all frequencies non-negative. The frequencies for the states are then *h*_*fast*_ *f*_0_ for the state ‘fast absent’, *h*_*fast*_ *f*_1_ for ‘fast present’, *h*_*slow*_ *f*_0_ for ‘slow absent’ and *h*_*slow*_ *f*_1_ for ‘slow present’. The rate matrix is multiplied with the frequencies over states and resulting matrix is exponentiated to obtain the matrix of transition probabilities. If *h*_*fast*_ = *h*_*slow*_ = 0.5 this is a time reversible process, but otherwise it is not, though a correction can be applied to make it reversible^1^.

Note there are 2 degrees of freedom for frequencies, one for visible and one for hidden frequencies, so together with switch rate *s* and mutation rate α this makes 4 degrees of freedom, as we shall see, that is one more than the pseudo Dollo model.

Since the covarion model allows multiple birth events in the tree, it can cater for non-Dollo like characters, and tends to fit well when there is some borrowing present in cognate data.

### 2.3 Stochastic Dollo model

The stochastic Dollo model [1, 15] is inspired by Dollo’s law of irreversibility, a hypothesis proposed by Louis Dollo in 1893 [5]. It states that oncea feature has been lost it will not be obtained again. This nicely fits with the definition of cognates, which are word forms with a common ancestry, obtained once at some point in history, and possibly lost many times, but not gained multiple times. The stochastic Dollo mode assumes each cognate can only arise once (with Poisson rate λ) but can be lost multiple times with death rate *µ*. Once the cognate is lost, it cannot arise again. The infinitesimal rate matrix of this process is

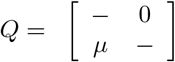

In general, the assumption of the stochastic Dollo model turn out to be too restrictive for cognate data as collected by linguists. Part of the reason is that horizontal transfer (that is borrowing of terms between languages in contact) results in more than one birth event of the cognate. Say language A borrows from B, but A and B are phylogenetically distant. For the stochastic Dollo model to fit such a situation, it has to assume a birth at or before the most recent common ancestor of A and B, which can be quite high in the tree, and may require a large number of losses of the cognate, resulting in a very low probability of such event. The same patterns are present in cognate data when there is semantic shift or coding errors. The covarion model tends to fare much better in such situations, since it allows multiple births of the cognate.

### 2.4 Pseudo Dollo model

Given the definition of cognates, it seems natural to assume that when a cognate becomes present, it can be lost, but is unlikely to be regained in accordance with Dollo’s law of irreversibility. However, there does not seem to be a reason to assume the restriction of the stochastic Dollo model that there can be only a single birth event of the cognate at some point in the tree. The pseudo Dollo model allows multiple births, but once a cognate is gained, it only allows the cognate to be lost once, never to be gained again.

The pseudo Dollo model is a CTMC model with three states: initial, present and removed. There are two rate parameters, the birth rate λ and death rate *μ* and the rate matrix Q for the CTMC has the following form

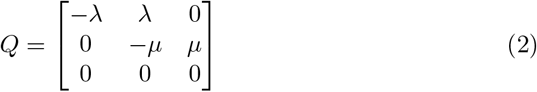

Note that this process (like the stochastic Dollo model) is not time reversible; once transitioned from the initial to the present state, the initial state cannot be entered any more, and once in the removed state there is no way to ape frescom it.

Let *T* be a rooted tree over a set of *n* taxa *x*_1_,…,*x*_*n*_. Internal nodes *x*_*n*+1_,… *x*_2*n*−1_ of the tree are numbered *n* + 1,…, 2*n* − 1. By convention, the highest numbered node (2*n* − 1) is the root and we use *π*_*i*_ to denote the index of the parent of node *i*. With each node *x*_*i*_ is associated a character *a*_*i*_, which is the ‘present’ state if the feature (say a cognate) is present, but if the feature is not present it is both ‘initial’ and ‘removed2019, so (like with the covarion model) we use ambiguity to encode characters. Then the likelihood of observing the set of tip characters **a**_**1**_…**a**_**n**_, given a tree *T* given, birth and death parameters λ and *μ* and other parameters *θ* (like branch length, etc.) is

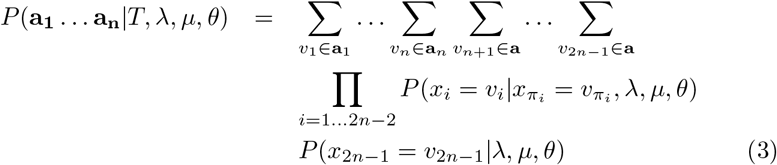

where the branch probabilities

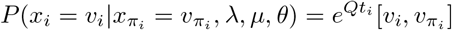

Here, *Q* is the rate matrix from Eq. (2). This is the probability of going from character 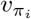 to *v*_*i*_ over time *t*_*i*_ according to the CTMC model. Note *t*_*i*_ is typically the branch length, but when using clock models(like the uncorrelated relaxed clock, see [6] forotherclock models) there may be a rate multiplier involved as well. Further,

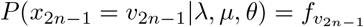

where *f*_*initial*_, *f*_*present*_ and *f*_*removed*_ are the so called frequency parameters at the base of the tree. Since they sum to one (*f*_*initial*_ + *f*_*present*_ + *f*_*removed*_ = 1 and all frequencies non-negative), there are only two degrees of freedom introduced. Eq. (3) looks very demanding, but fortunately it can be calculated efficiently using Felsenstein’s pruning algorithm [9], which allows it to be calculated in *O*(*n*) time.

Since typically a clock model is assumed, which provides a multiplier for the branch lengths, we cannot estimate both λ and *μ* as well as the clock rate; raising the clock rate and reducing both λ and *μ* results in the same fit, so we have an identifiability issue. By fixing λ to 1, this problem is solved, leaving just 3 degrees of freedom for the psuedo Dollo model: one for *μ* and two for frequencies.

We assume a Dirichlet(1,1,1) prior over the frequencies and an exponential with mean 1 on the death rate, unless specified otherwise.

In practice, it is faster to compute Eq. (3) using an extra dummy state, since current computer processors are optimised to perform operations on blocks of 4. So, instead of using the Q-matrix from (2), we use a four state matrix

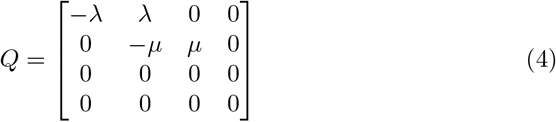

and set the frequencies for the dummy state to zero. The character encoding is like before, so the number of degrees of freedom of the model does not change.

#### Typical usage

The pseudo Dollo model can be used as alternative to the stochastic Dollo model.

We envision that the pseudo Dollo model will be used for cognate data according to the scheme of Chang et al. [4] as follows: for each meaning class, a separate mutation rate is introduced, which is a multiplier for the branch length that is independent from the clock rate. The mean of mutation rates over meaning classes (weighted with the number of cognates in the meaning class) is kept to 1 to ensure identifiability. Each of the meaning classes has ascertainment correction conditioned on the languages having data for the particular meaning class being absent - in effect conditioning on the fact at least some of the languages contain the cognate, otherwise the cognate would not have been added to the dataset. The meaning classes can share the pseudo Dollo model, so there will be just 3 parameters to estimate for the Dollo model.

When there are few cognates per meaning class, it may be better to join the cognates in a single partition and use a single mutation rate instead of one rate per meaning class. In this situation, the pseudo Dollo model can benefit from gamma rate heterogeneity across sites [19].

### 2.5 Variations on the pseudo Dollo model

We introduce two generalisations of the pseudo Dollo model. The first one allows for fast and slow modes like in the covarion model, which we call the pseudo Dollo covarion model. The second variation of the pseudo Dollo model allows distinguishing different present states, which we call the multi state pseudo Dollo model.

#### Pseudo Dollo Covarion model

The success of the covarion model in capturing cognate evolution suggests some cognates can be stable (either absent or present) for a long period of time alternated with rapid evolution. This feature of cognate evolution is not captured by the pseudo Dollo model; it assumes constant rate of evolution along all branches in the tree. Just like the covarion model expands the state space of the binary CTCM model with an extra state, we can expand the statespace of the pseudo Dollo model with one extra state for each of the tree states to capture the switch between fast and slow evolution. This would yield a fast and a slow absent state,a fast and a slow present state and a fast and slow removed state. Since it will be impossible to escape the removed state, the fast and slow removed states can be collapsed into a single state. The rate matrix for the pseudo Dollo (PD) covarion model with states *fastabsent*, *fast present*, *removed*, *slow absent*, *slow present* is then

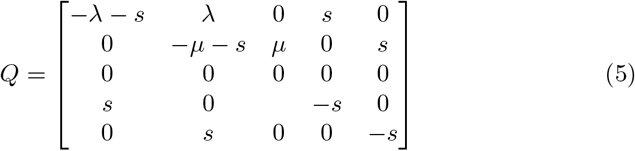

**Figure 1:**
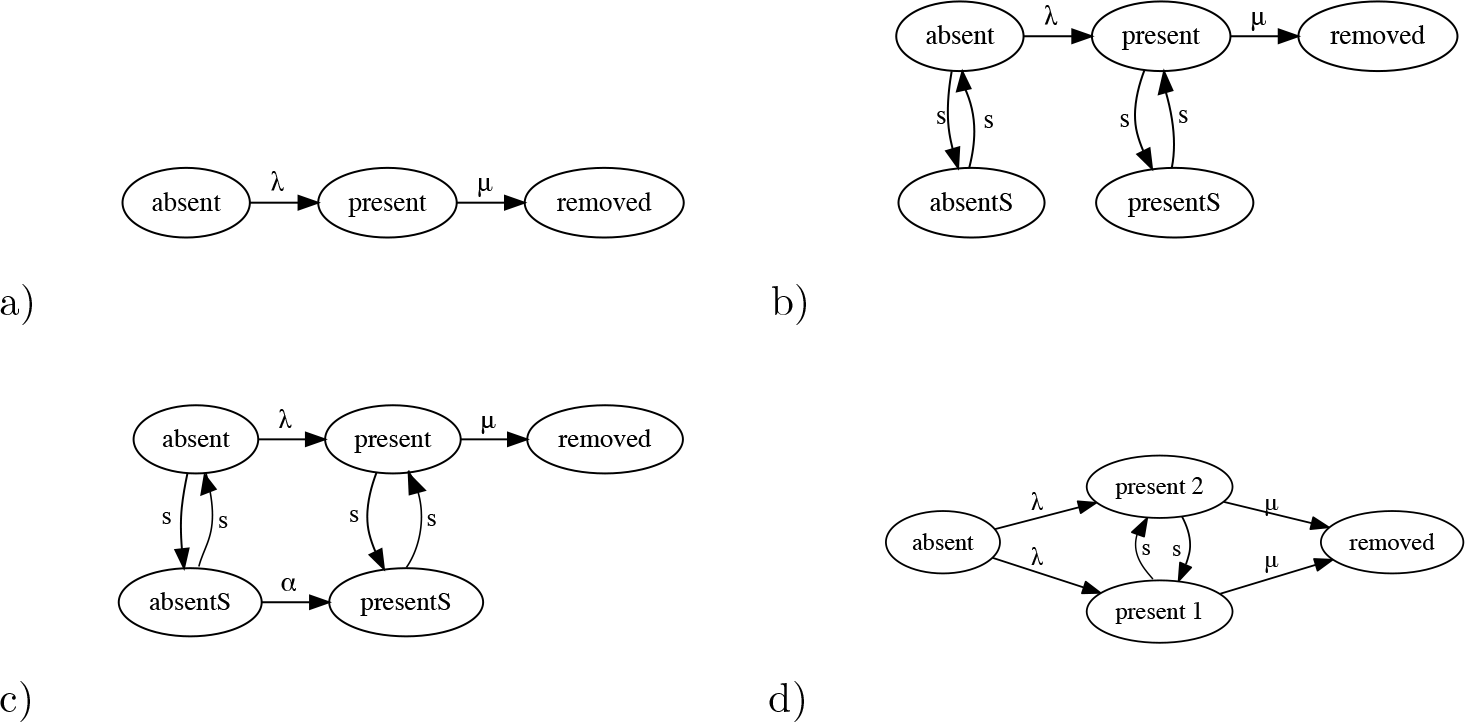
Diagrams for (a) pseudo Dollo, (b) PD-covarion, (c) PD-covarion B, and (d) multistate pseudo Dollo models with all rate parameters.

Like the pseudo Dollo model, we have frequencies for absent, present and removed states. Like for the covarion model, this is extended by frequencies for fast and slow states, easiest kept to 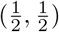 like in the reversible covarion model,giving frequencies 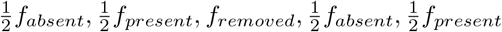.

Unlike Tuffley and Steel’s covarion model, which does not allow a slow 0 to change into a slow 1 or vice versa, the BEAST covarion model allows such changes with a small rate *α*. In practice, this can lead to a large increase in the fit of the model. The PD-covarion model can be allowed a change from slow 0 to slow 1 with rate small rate *α* (*α* ≪ λ)giving rise to the following model at the cost of an extra parameter

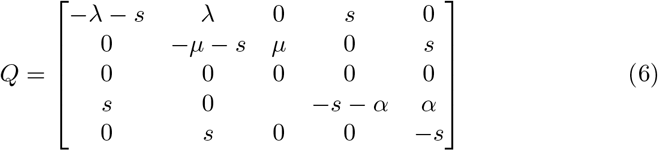

#### Multi state pseudo Dollo model

One way to encode cognates is by taking semantic shift in account. For example,when there are three meanings for a cognate, instead of encoding them in binary form where 0 means absence and 1 presence of the cognate, we can distinguish presence of the cognate with three different values, say 1, 2 and 3. by allowing the present state to have multiple values that are allowed to transition amongst themselves. For example, when there are 3 possible states recognised as being present, *p*_1_, *p*_2_, *p*_3_, we have states absent, *p*_1_, *p*_2_, *p*_3_, *removed* with ratematrix

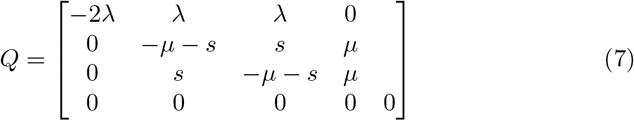

The stochastic Dollo model was generalised to include more than two states in [1]. A similar generalisation is possible for the pseudo Dollo covarion model where the *present* state is split into multiple states, each with their own fast and slow states, and allowing rates among fast present states.

## 3 Results

We performed a simulation study to ensure parameters can be reliably estimated from simulated data. Since BEAST normalises the rate matrix so that branch lengths can be interpreted in expected number of mutations per site, one parameter can be chosen to be fixed, and we set the birth rate λ to 1. A death rate and frequencies was sampled from the prior (*μ* from exponential with mean 1 truncated to the range 0 to 1, and frequencies from Dirichlet(5,2,1)). The tree was sampled over 10 taxa using a Yule prior with birth rate sampled from a log normal with mean 4 (in real space) and variance of 0.05. We used a strict clock model to simulate sequence data of 1000 characters using the sequence simulator facility in BEAST [2]. Then we ran an MCMC analysis in BEAST under the same model, with a Yule tree prior co-estimating the phylogeny and model parameters for 50 million samples, thus guaranteeing convergence. We repeated the procedure a hundred times, and expected the model parameters used to simulate the data is in the 95% highest probability density (HPD) interval at about 95% of the runs. Table 1 summarises results, and shows that indeed approximately 95% of the time the true value were contained in the 95% HPD interval.

**Table 1.** Results of simulation study for pseudo Dollo model. The number indicates how many of the 100 simulated data sets resulted in inferences where the true value (used to simulate the data) was contained in the 95% highest probability density (HPD) interval.

| parameter | coverage |
| --- | --- |
| $\mu$ | 96 |
| $f_{absent}$ | 92 |
| $f_{present}$ | 96 |
| $f_{removed}$ | 93 |
| Tree height | 95 |
| Yule birth rate | 94 |

We compared performance of the pseudo Dollo model with that of the 2 state CTMC and covarion model on three different datasets: Indo-European cognate data from the IELex database [3] with cognate data updated in 2015, and Transeurasian data among 5 language families. The first dataset contains many entries per meaning class, but the latter doesn't, so for the first analyses we used the scheme of Chang et al. [4] allowing one mutation rate per meaning class, but for the latter we used a single partition analysis.

For the Indo-European analysis, we used a rather sophisticated sampled ancestor analysis [10] allowing ancient languages to become ancestral to later languages, including modern day languages. An uncorrelated relaxed clock [7] was used as clock model and calibrations similar to the ones in [3] were used. In previous studies [3] it was shown that the covarion model outperforms the CTMC and stochastic Dollo model by a considerable margin on similar data. There is no reason to believe this changes with the corrections that were made to the data, but since these analyses take quite long to run we only considered the covarion and pseudo Dollo model. The marginal likelihoods for covarion and pseudo Dollo did not even overlap. The mean likelihood was −51202 and −40212 for these models respectively, indicating a much better fit for the pseudo Dollo model.

The Transeurasian data consists of cognate data over 50 languages spread among 5 language families: Japonic, Koreanic, Mongolic, Tungusic and Turkic. A detailed description of the data can be found in [16]. For the trans-Eurasian data, we performed a uncorrelated relaxed clock [8] analysis with Yule prior.

Table 2 shows marginal likelihood estimates obtained using nested sampling [14], which has as added benefit that it estimates both log marginal likelihood and standard deviation of the estimate. The pseudo Dollo model outperforms the covarion model, two state CTMC and stochastic Dollo models. with Bayes factors overwhelmingly [13] favouring it. Adding rate heterogeneity to the pseudo Dollo model improves the marginal likelihood. But the model fit for the pseudo Dollo covarion model is even better than any of the other models. The stochastic Dollo model performs especially poor compared to the other models.

**Table 2.** Log marginal likelihood (ML) estimates (standard deviation in brackets) for Transeurasian data for various substitution models using the relaxed clock. Higher ML estimates indicate better model fit. Difference of ML estimates give the log Bayes factor(BF). This indicates that the pseudo Dollo covarion model gives BF≫100 with respect to any of the other models, and is the best for for the data.

| Substitution model | Log Marginal likelihood |
| --- | --- |
| Covarion model | -1090.8 (0.99) |
| Two state CTMC model | -1106.5 (0.96) |
| Two state CTMC, 4 gamma model | -1098.7 (1.05) |
| Stochastic Dollo model | -1447.1 (1.74) |
| Pseudo Dollo model | -1028.1 (0.92) |
| Pseudo Dollo, 4 gamma model | -1020.1 (0.93) |
| Pseudo Dollo covarion model | -1015.5 (0.93) |

## 4 Conclusion

We showed that the pseudo Dollo has a comparable or even better fir than other models commonly used in cognate evolution. In other words, the pseudo Dollo model has the potential to explain the data equally well or better than other popular models. In particular, it performs much better than the stochastic Dollo model, hence provides a good alternative to the other models for modelling and explaining the data.

The main difference between the stochastic Dollo and the pseudo Dollo model is that the stochastic Dollo model does not allow more than one birth event, but the pseudo Dollo model does. As a consequence, when a cognate is borrowed from a distant language, the stochastic Dollo model explains this by having a birth event before the most recent common ancestor (MRCA) of the languages involved, and for any of the descending clades that does not have the cognate present a death event. When the clade borrowed from is distant, a lot of death events are required, each death event making this scenario less likely. The pseudo Dollo model on the other hand requires just two birth events, one each before the clades containing the cognate, and requires less death events. On balance, the pseudo Dollo model explains the data better when a borrowing event takes place from a distant branch.

Likewise, when there are errors in coding the data, the stochastic Dollo model requires a birth event before the MRCA of all taxa containing the cognate, including the miscoded one, and thus includes possibly a large number of death events. The pseudo Dollo model on the other hand only requires two birth events; one before the MRCA of the actual cognate set, and one before the miscoded taxon. In this case, the pseudo Dollo model fits the data better. In addition, the covarion model allows multiple birth events, so it can deal with miscoded cognates in a similar way.

The pseudo Dollo and PD-covarion models are implemented in Babel, a package for BEAST [2], and a BEAUti template is available for setting up an analysis using a graphical user interface. The package is licensed under LGPL, and source code is available from https://github.com/rbouckaert/Babel.

## Acknowledgements

This research was funded by Marsden grant (UOA1308) (http://www.royalsociety.org.nz/programmes/funds/marsden/awards/2013-awards/), a Rutherford fellowship (http://www.royalsociety.org.nz/programmes/funds/rutherford-discovery/)from the Royal Society of New Zealand awarded to Prof. Alexei Drummond. It was also funded by the Max Planck Institute for the Science of Human History.

## Footnotes

1 M. Pagel personal communication.

